# Absolute quantification of TCA cycle intermediates in mouse ocular tissues reveals distinct tissue- and sex-specific mitochondrial metabolism

**DOI:** 10.1101/2025.09.25.678562

**Authors:** Cloe Ratliff, David Hansman, Tuan Ngo, Yinxiao Xiang, Artjola Puja, Mark Eminhizer, Jinyu Lu, Isabella Mascari, Diana Alabdallat, Jianhai Du

## Abstract

**Objective:** Mitochondrial tricarboxylic acid (TCA) cycle is central to energy production and redox balance in the eye, which must sustain high metabolic activity to support vision. Retinal neurons, the retinal pigment epithelium (RPE), cornea, and lens each have distinct physiological roles and metabolic demands, yet the absolute concentrations of key TCA intermediates and their variation by tissue, sex, and time of day are not well-defined.

**Methods:** Targeted gas chromatography–mass spectrometry was employed to quantify the absolute concentrations of TCA cycle metabolites in mouse ocular tissues collected at 10 AM and 2 PM to capture diurnal variations. Key metabolite ratios were subsequently calculated to provide insight into TCA cycle dynamics across eye tissues.

**Results:** The retina showed the highest concentrations of TCA metabolites among all ocular tissues, particularly succinate, citrate, and malate, consistent with its high energy demands. The RPE/choroid demonstrated well-balanced intermediates with the highest α-ketoglutarate (α-KG)/Isocitrate ratio, reflecting its efficient mitochondrial oxidation and reductive carboxylation. Corneal metabolism was featured by dominant malate, especially in females, suggesting a metabolic adaptation for redox regulation and oxidative stress defense. The lens had uniformly low metabolite levels except for succinate, indicating minimal mitochondrial activity under physiologically low oxygen conditions. Notably, both the cornea and lens showed significant sex-dependent and diurnal variations in TCA cycle intermediates.

**Conclusion:** This study demonstrates distinct tissue-specific mitochondrial metabolism in the eye, reflecting the unique functional and biochemical demands of each tissue. These metabolic signatures may underlie their susceptibility to mitochondrial dysfunction in various ocular diseases.

## 1. Introduction

The TCA cycle lies at the heart of mitochondrial metabolism, a hub that integrates central carbon metabolism with energy production, biosynthesis, and redox functions^1^ (Figure 1A). It oxidizes carbon fuels, including glucose, amino acids, fatty acids and ketone bodies, powering the electron transport chain (ETC) to drive ATP synthesis^2,3^. TCA intermediates also provide key precursors for the biosynthesis of lipids, amino acids, and nucleotides. For example, citrate provides cytosolic acetyl-CoA for fatty acid, cholesterol, and phospholipid synthesis^4^, while α KG provides carbons for the synthesis of glutamate, proline, glutamine, and aspartate^5^. Moreover, TCA metabolites such as succinate, fumarate, and malate regulate diverse cellular processes, including immune modulation, antioxidant defense, and epigenetic regulation^1^.

**Figure 1.**
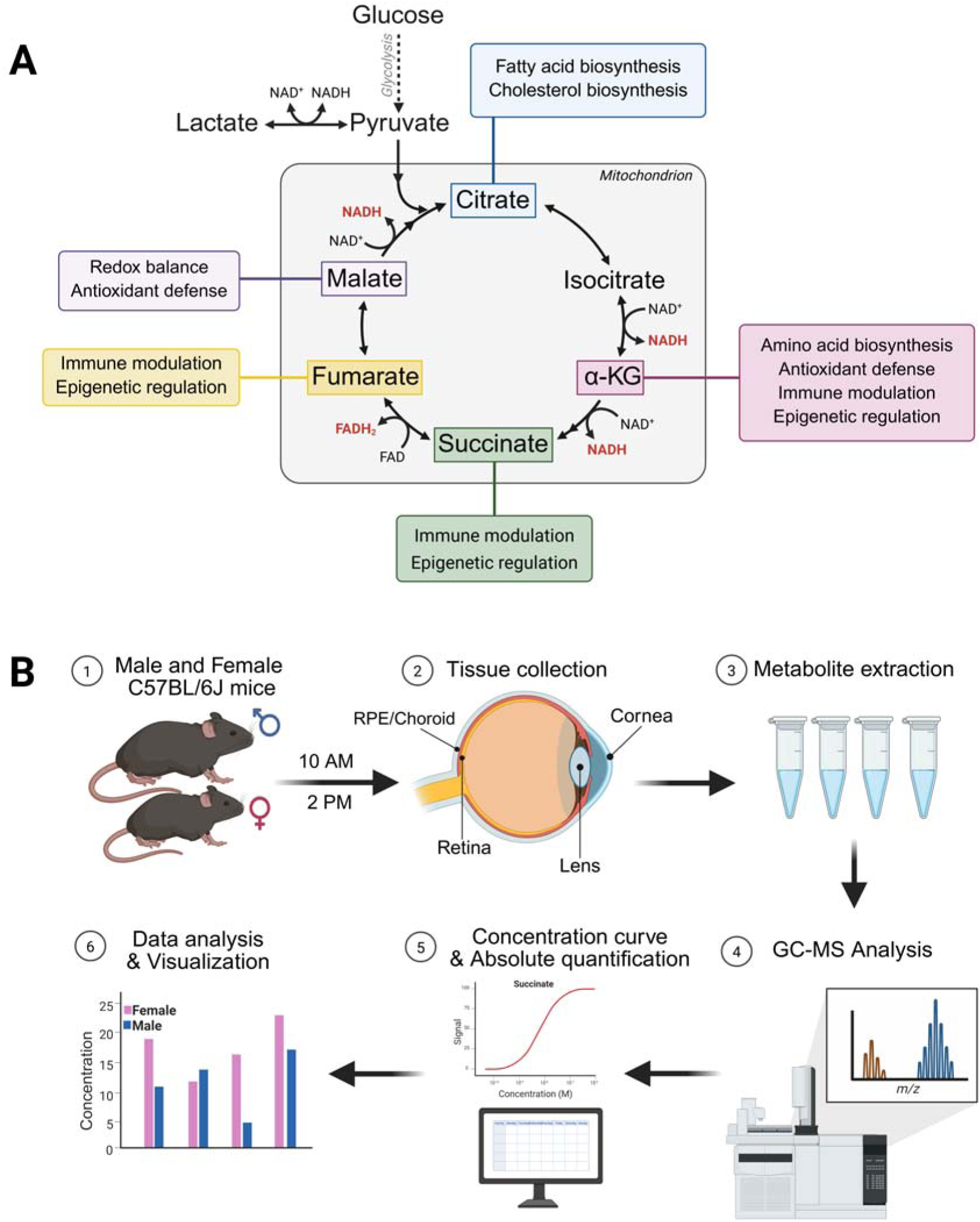
TCA cycle overview and study methodology. (A) The TCA cycle is a central metabolic hub that integrates energy production, biosynthesis, and redox functions. It supports ATP production through the generation of the reducing equivalents NADH and FADH (shown in red). TCA cycle intermediates are precursors for the synthesis of lipids, amino acids, and other metabolites. Metabolites such as -KG, succinate, fumarate, and malate have diverse functions regulating redox balance, antioxidant defense, epigenetics, and immune functions. (B) (1) TCA cycle metabolites were analyzed in male and female C57BL/6J mouse eye tissues collected at 10 AM and 2 PM. (2) The retina, RPE/choroid, lens, and cornea were removed and snap frozen for later analysis. Tissue metabolites were extracted (3) and analyzed using gas chromatography– mass spectrometry (GC-MS) (4), generating spectra of relative abundance versus mass-to-charge ratio (m/z). (5) Data quantification identified metabolites of interest and external standards were prepared for absolute quantification. (6) Statistical analyses revealed sex-dependent metabolic differences across ocular tissues.

The retina is the most metabolically demanding tissue in the body^6^, and relies on both aerobic glycolysis and oxidative phosphorylation (OXPHOS) to support its profound bioenergetic and biosynthetic demands^7,8^. This unusual metabolic program is driven largely by the photoreceptors, which must expend energy to control ion fluxes associated with phototransduction, continually synthesize components of their outer segments, as well as other energetic burdens that support their unique physiology^9–11^. The RPE, a polarized monolayer of cells overlying the retina, provides a cellular barrier that regulates nutrient transport between the outer retina and circulation^12^. We previously reported that the healthy RPE has extremely high metabolic flexibility, utilizing more than 50 nutrients in its mitochondria^13^. RPE cells have abundant mitochondria that possess a high capacity to process nutrients and recycle “waste products,” such as lactate and phagocytosed outer segments from circulation and the outer retina, while synthesizing and exporting nutrients to support the outer retina^13–18^. Mitochondrial defects and impaired OXPHOS in the RPE and retina are associated with many retinal diseases, including age-related macular degeneration (AMD)^19–26^, diabetic retinopathy^23,24,26^, retinitis pigmentosa^27–29^, and glaucoma^23,30^.

The cornea consists of a thick, transparent stromal layer of regularly arranged collagen fibers, lined by an epithelial layer on the anterior and a monolayer of endothelial cells on the posterior side^31,32^. While the whole cornea predominantly relies on glycolytic metabolism^33,34^, corneal endothelial cells contain a dense population of mitochondria, using both glucose and glutamine as carbon sources for the TCA cycle^35^. TCA cycle-derived ATP is critical to fuel the ion and lactate transport pumps in corneal endothelium that maintain stromal hydration and transparency^35,36^. While the corneal epithelium is largely glycolytic^37,38^, it relies on mitochondrial metabolism for redox balance, differentiation, and wound healing^39–42^. The turnover rate of the corneal stroma extracellular matrix (ECM) is extremely slow, making it vulnerable to oxidative damage^43,44^, while TCA metabolites such as malate and α-KG are key defenders against oxidative stress by modulating NADH, NADPH, and glutathione production and by increasing antioxidant enzyme activity^45,46^. While most of the lens is composed of mature fiber cells that lose their mitochondria during differentiation to rely on glycolysis, the monolayer of epithelial cells overlying the anterior lens is rich in small mitochondria^47–49^. The TCA cycle in the lens epithelium provides ATP production, redox regulation, and defense against oxidative stress^50,51^. These functions are crucial for maintaining lens transparency and cellular integrity. Mitochondrial function is critical to both the cornea and lens, and its dysfunction is associated with a number of outer eye diseases, including Fuchs endothelial corneal dystrophy, congenital hereditary endothelial dystrophy, and age-related cataracts^36,42,52–56^.

Sexual dimorphisms have been reported in visual physiology, function, metabolism, and susceptibility to eye diseases^57–60^. Substantial differences between male and female mice have been identified in the RPE and retinal proteomes^61^. Moreover, we recently reported that the mouse RPE, retina, and lens show sex-specific metabolic responses to fasting, influencing amino acid, nucleotide, lipid, and sugar metabolites^62^.

Despite the importance of mitochondrial metabolism in eye diseases, the concentrations of mitochondrial substrates in healthy ocular tissues, and how they differ between sexes, remain unclear. In this study, we employ targeted metabolomics to quantify the absolute concentrations of TCA cycle intermediates in the retina, RPE/choroid, cornea, and lens of male and female mice at 10 AM and 2 PM. We identify strikingly different concentrations and ratios of mitochondrial intermediates in ocular tissues, revealing their unique mitochondrial metabolism. These findings provide novel insight into how mitochondrial metabolism is biochemically adapted to support specific functions in ocular tissues, suggesting unique vulnerabilities to mitochondrial dysfunction in eye diseases.

## 2. Material and methods

### 2.1. Animals

Eight-week-old C57BL/6J wild-type mice (6 male, 6 female) were purchased from Jackson Laboratory (Detailed in Table S1). All mice were maintained on ad libitum feeding and acclimated to the facility environment for one week prior to experimental procedures. All mouse experiments were performed in accordance with guidelines from the National Institutes of Health and ARVO Statement for the Use of Animals in Ophthalmic and Vision Research, and the protocols were approved by the Institutional Animal Care and Use Committee of West Virginia University.

### 2.2. Tissue Collection

Six mice (3 male, 3 female) were used for each of the 10 AM and 2 PM time point. The mice were euthanized via quick cervical dislocation, and eyes were enucleated with care to remove as much excess connective tissue as possible. The eyes were then dissected to isolate the retina, RPE/choroid, cornea, and lens as reported previously^63,64^. Tissues were then snap-frozen in liquid nitrogen and stored at –80 for further processing.

### 2.3. Metabolite Extraction and Protein Assay

Metabolites were extracted from the retina, RPE/choroid, cornea, and lens using 80% cold methanol containing 1 mM norvaline as an internal standard, as previously reported^65^. After extraction, the mixtures were centrifuged, and a 10 µL aliquot of each supernatant was dried, derivatized, and analyzed by gas chromatography–mass spectrometry (GC–MS) (Agilent 7890B/5977B instrument) in selected ion monitoring mode, as previously described^66^. The TCA cycle intermediates citrate, isocitrate, α-KG, succinate, fumarate, and malate, as well as lactate and pyruvate, were targeted for analysis (Table S2). Proteins in the residual pellet were extracted with 1 M NaOH and incubated overnight in a thermomixer at 37 °C to solubilize proteins. Protein concentrations were quantified using the Pierce™ BCA Protein Assay Kit (Table S1) and were used for normalization of metabolite concentrations^67^.

### 2.4. Metabolite Quantification

Absolute quantification was performed using external calibration curves containing internal standards for each targeted metabolite. Calibration standards spanning the physiological range were analyzed under the same chromatographic conditions (Table S3). Linear regression of peak area versus concentration (R² > 0.99) was used to calculate metabolite levels in each sample. Values were corrected for reconstitution volume and normalized to protein concentration, as determined by the protein assay, and are reported as nanomoles per milligram of protein (Table S4). All raw mass spectrometry data have been deposited to MassIVE (dataset identifier: MSV000098916).

### 2.5. Statistical Analysis

Data are presented as the mean ± standard deviation (SD). Statistical analysis was performed using Student’s unpaired, two-tailed t-tests or a one-way ANOVA followed by the Bonferroni post hoc test, for multiple comparisons using GraphPad Prism 10.0. Metabolite ratios were calculated for each metabolite pair by dividing the mean value of the denominator metabolite (across all samples in the group) by the individual values of the numerator metabolite for each metabolite pair. A *P*-value of < 0.05 was considered statistically significant.

## 3. Results

### 3.1. The retina has abundant TCA intermediates with succinate accumulation

To quantify TCA intermediates across different ocular tissues, we analyzed metabolites from freshly isolated retina, RPE/choroid, cornea, and lens tissues from male and female mice, collected at 10 AM and 2 PM, using GC-MS (Figure 1B). Our analysis focused on key TCA cycle intermediates, including citrate, isocitrate, α-KG, succinate, fumarate, and malate, as well as the glycolytic end products, pyruvate and lactate.

In the retina, no significant differences in metabolite levels were observed between male and female mice at either 10 AM or 2 PM (Figure 2A, B). Moreover, metabolite concentrations remained largely stable between both time points (Figure S1), with lactate levels far exceeding those of TCA metabolites by >200-fold (Figure 2A, B), consistent with the Warburg effect in the retina^8^. Among TCA metabolites, succinate was consistently the most abundant in both sexes, followed by malate, citrate, and isocitrate (Figure 2C-F). In contrast, fumarate and α-KG were the least abundant, with concentrations approximately 10-fold lower than succinate. This high succinate-to-fumarate ratio suggests reverse succinate dehydrogenase (SDH) activity for electron transport under hypoxic conditions^68,69^. The abundant citrate and malate may facilitate lipid biosynthesis and malate–aspartate shuttle (MAS) activity^70,71^. These findings suggest that the retina adapts its TCA cycle to maintain energy production and biosynthetic capacity in a low-oxygen environment.

**Figure 2.**
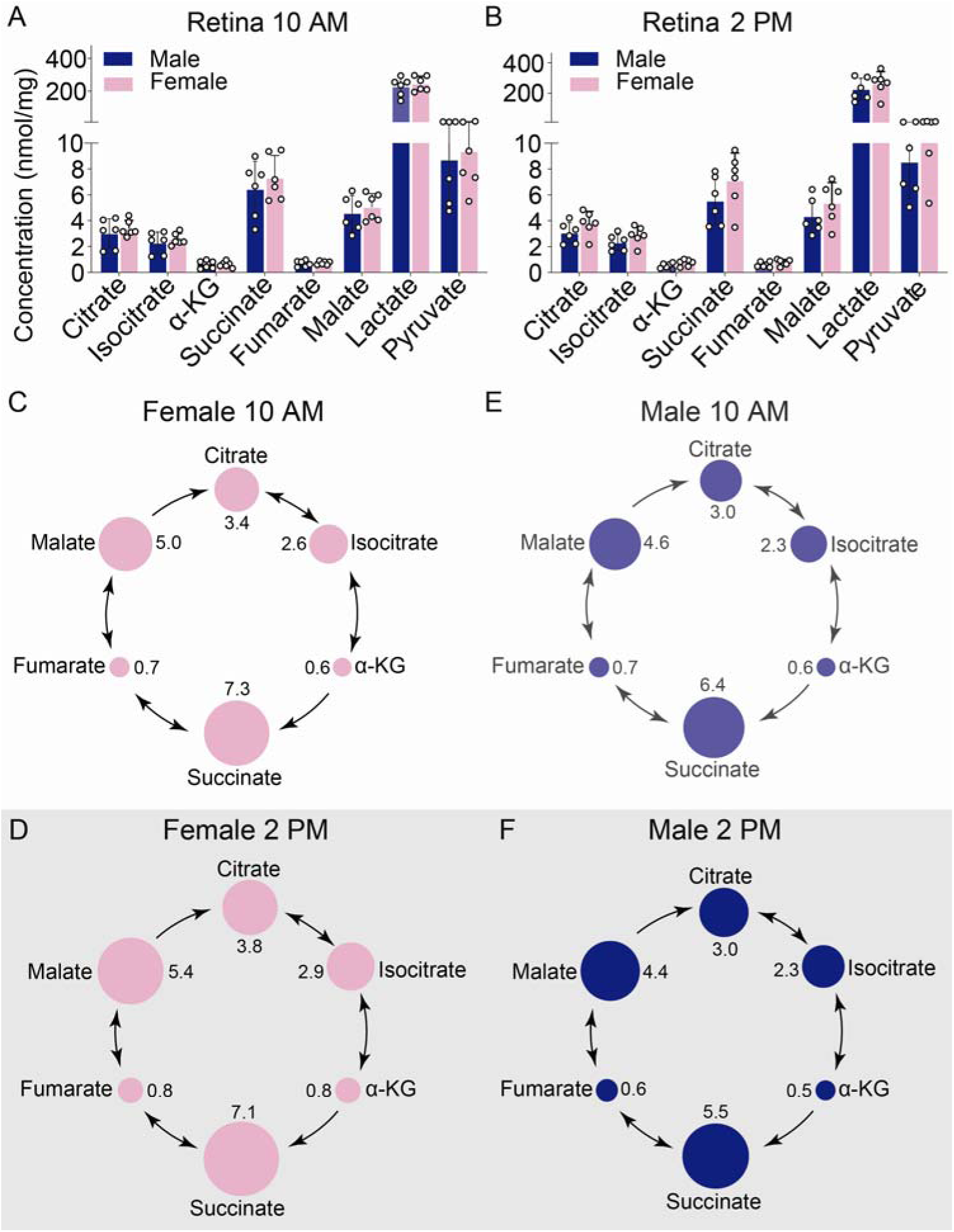
Quantification of retinal TCA cycle metabolites in male and female mice at 10 AM and 2 PM. Lactate, pyruvate, and TCA metabolite concentrations in male and female mouse retinas collected at (A) 10 AM and (B) 2 PM. No statistical significance (p<0.05) between male and female samples were detected using an unpaired, two-tailed Student’s t-test. Visualization of retinal TCA cycle metabolite abundances in female mice at 10 AM (C) and 2 PM (D), and male mice at 10 AM (E) and 2 PM (F). Circle sizes represent metabolite abundance and are scaled consistently across figures. Numbers alongside circles are abundance values normalized to protein concentration.

### 3.2. The RPE/choroid maintains well-balanced TCA intermediates

In the RPE/choroid, levels of TCA cycle intermediates showed no significant differences between males and females at 10 AM (Figure 3A). However, at 2 PM, succinate concentrations in male mice decreased to 0.4 nmol/mg, while levels in females remained stable at approximately 0.7 nmol/mg (Figure 3B). Statistically significant decreases in α-KG and fumarate were also observed in male mice between 10 AM and 2 PM (Figure S2). Otherwise, metabolite concentrations were remarkably consistent across sexes and time points, and the relative distribution of TCA intermediates showed minimal variation (Figure 3A-F). As in the retina, lactate and pyruvate were the most abundant metabolites (Figure 3A, B). Among TCA cycle metabolites, malate and succinate were present at the highest levels, while citrate, isocitrate, and α-KG were consistently the least abundant, independent of sex or timepoint (Figure 3C-F). Overall, the profile of the RPE/choroid showed well-balanced TCA metabolite levels, except for moderately elevated malate and succinate.

**Figure 3.**
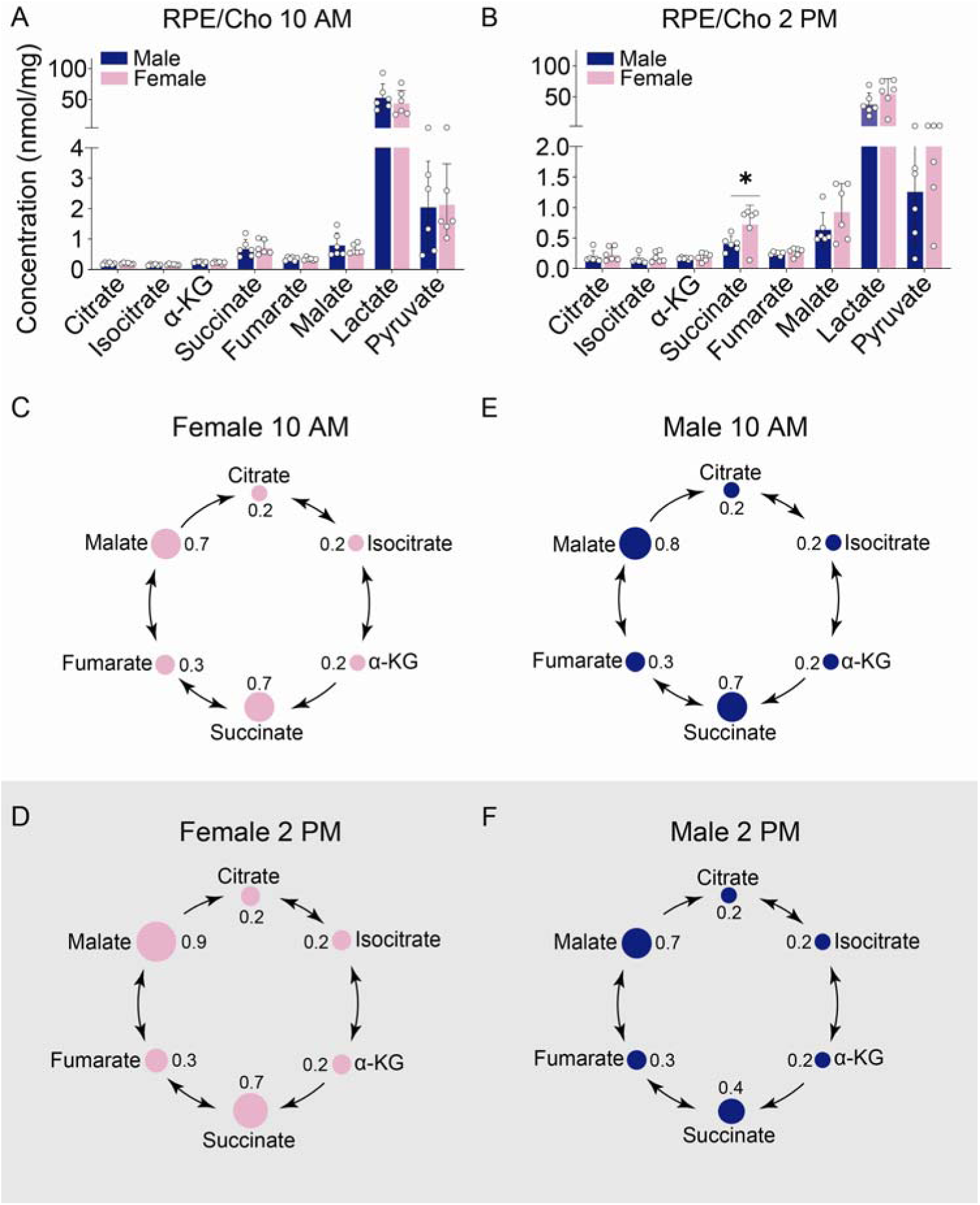
Quantification of RPE/choroid TCA cycle metabolites in male and female mice at 10 AM and 2 PM. Lactate, pyruvate, and TCA metabolite concentrations in male and female mouse retinas collected at (A) 10 AM and (B) 2 PM. Statistical significance: *P<0.05 between male and female samples. Visualization of RPE/choroid TCA cycle metabolite abundances in female mice at 10 AM (C) and 2 PM (D), and male mice at 10 AM (E) and 2 PM (F). Circle sizes represent metabolite abundance and are scaled consistently across figures. Numbers alongside circles are abundance values normalized to protein concentration.

### 3.3. The cornea has high malate with more pronounced sex differences in the morning

Compared to the retina and RPE/choroid, the cornea showed more significant sex differences in mitochondrial intermediates at 10 AM. Female corneas exhibited 2-to 3-fold higher levels of citrate, α-KG, fumarate, and malate compared to males (Figure 4A). In contrast, no significant sex differences appeared at 2 PM; however, metabolite abundances at this time point showed considerable variability within groups (Figure 4B). No metabolites in either sex showed significant differences between 10 AM and 2 PM (Figure S3). Lactate was the most abundant metabolite across all groups (Figure 4A, B). Among TCA intermediates, malate had the highest concentration, particularly in the morning (Figure 4C-F). Interestingly, there was a surge in succinate concentration of 2-4-fold at 2 PM, especially in the female cornea (Figure 4C-F). Taken together, these results suggest that corneal TCA metabolism varies in a sex-dependent manner, while maintaining consistently elevated levels of malate.

**Figure 4.**
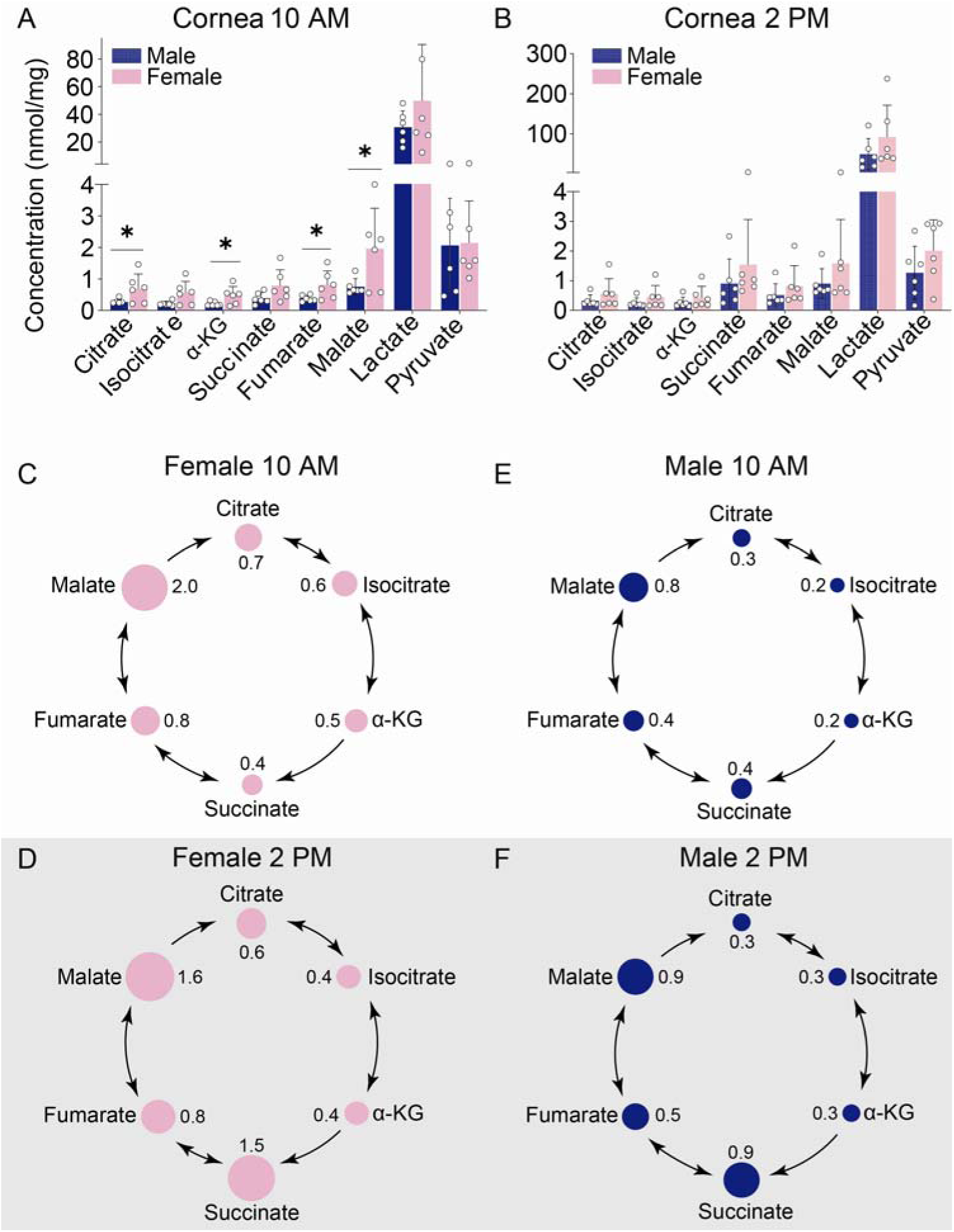
Quantification of corneal TCA cycle metabolites in male and female mice at 10 AM and 2 PM. Lactate, pyruvate, and TCA metabolite concentrations in male and female mouse corneas collected at (A) 10 AM and (B) 2 PM. *P<0.05 between male and female samples. Visualization of corneal TCA cycle metabolite abundances in female mice at 10 AM (C) and 2 PM (D), and male mice at 10 AM (E) and 2 PM (F). Circle sizes represent metabolite abundance and are scaled consistently across figures. Numbers alongside circles are abundance values normalized to protein concentration.

### 3.4. The lens has very low TCA intermediates, except for high succinate

In contrast to the cornea, sex-specific differences in lens metabolites were observed at 2 PM but not at 10 AM (Figure 5A). Concentrations of lactate, pyruvate, and all TCA cycle metabolites declined significantly in males compared to females (Figure 5B) and in males between 10 AM and 2 PM (Figure S4). As in other tissues, lactate was the most abundant metabolite in both male and female lenses at both timepoints (Figure 5A, B). Overall, TCA metabolite levels in the lens were very low, with the notable exception of succinate, which was by far the most abundant independent or variable (Figure 5C-F). In contrast, citrate, isocitrate, α-KG, and fumarate were present at minimal levels. In summary, the lens contained extremely low TCA metabolite levels overall, except for a high abundance of succinate, while these metabolites all declined in male mice by 2 PM.

**Figure 5.**
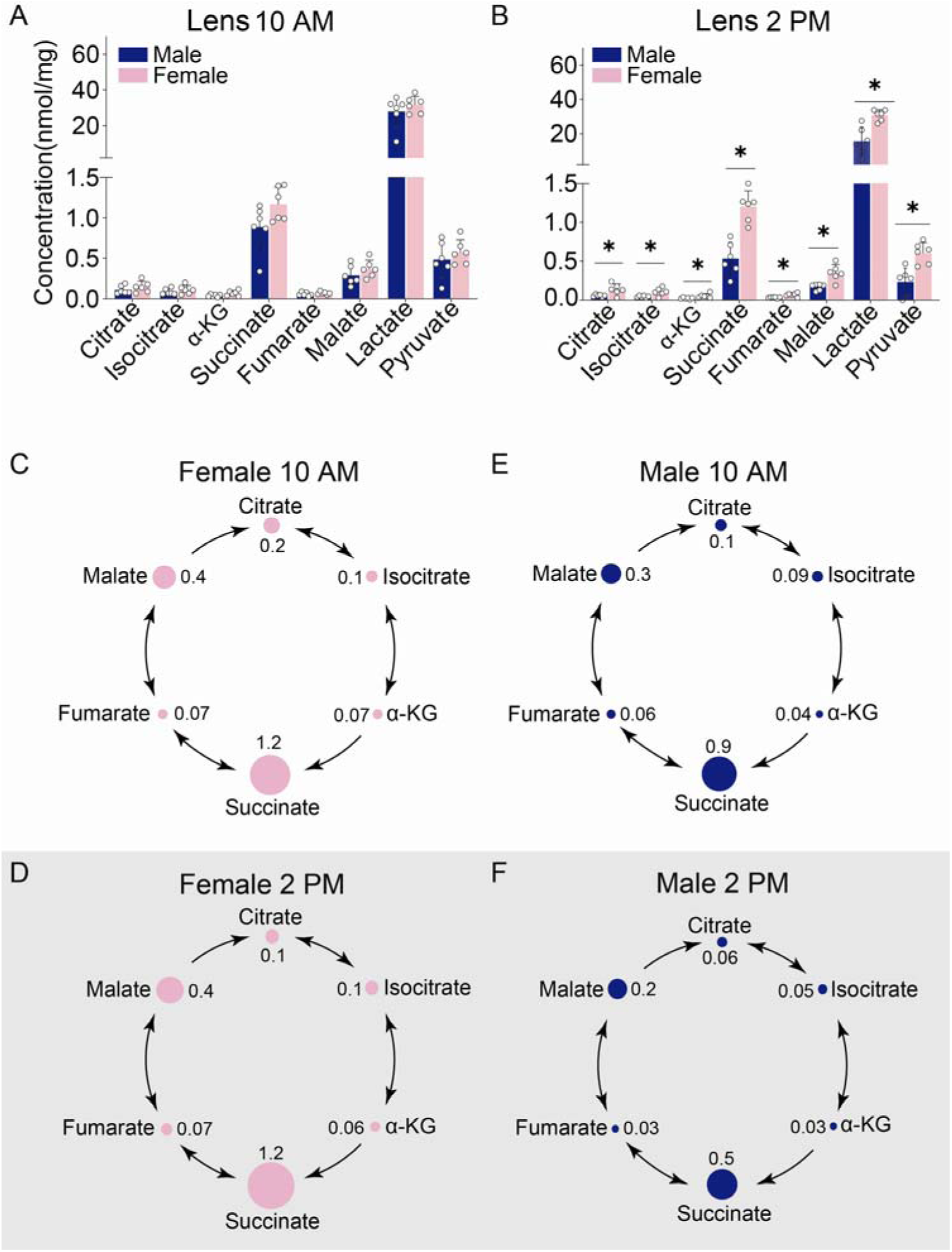
Quantification of lens TCA cycle metabolites in male and female mice at 10 AM and 2 PM. Lactate, pyruvate, and TCA metabolite concentrations in male and female mouse lenses collected at (A) 10 AM and (B) 2 PM. *P<0.05 between male and female samples. Visualization of lens TCA cycle metabolite abundances in female mice at 10 AM (C) and 2 PM (D), and male mice at 10 AM (E) and 2 PM (F). Circle sizes represent metabolite abundance and are scaled consistently across figures. Numbers alongside circles are abundance values normalized to protein concentration.

### 3.5. Metabolite ratios reveal tissue-specific TCA cycle activity

The relative levels of substrates and products are important indicators of TCA cycle enzyme activity. To further analyze TCA cycle dynamics across eye tissues, we calculated ratios of different intermediates including succinate/fumarate, isocitrate/citrate, α-KG/isocitrate, α-KG/succinate and citrate/α-KG (Figure 6A, B, Table S5). The lens and retina had the highest succinate/fumarate ratios, supporting their adaptation to use fumarate as an electron acceptor under limited oxygen^68,69^. Isocitrate/citrate was the most consistent ratio in all tissues, suggesting efficient aconitase activity throughout the eye. The α-KG/isocitrate ratio is directly related to reverse isocitrate dehydrogenase activity. As expected, the RPE/choroid had the highest α-KG/isocitrate ratio in both the morning and afternoon, consistent with our previous study showing that the RPE has a high capacity for reductive carboxylation^72^. The α-KG/succinate ratio is characteristic of high α-KG dehydrogenase activity, as well as non-TCA cycle enzymes including dioxygenases such as DNA, RNA, and histone demethylases, and prolyl hydroxylases^73,74^. Remarkably, the highest α-KG/succinate ratio was found in the cornea in the morning, and the RPE/choroid in the afternoon, with opposite sex differences (Figure 6). The malate/fumarate ratio reflects the cellular redox state and MAS activity^70^. The retina had the highest ratio, followed by the lens, suggesting a high demand for redox regulation. The citrate/α-KG ratio represents the metabolic decision point between energy production and biosynthetic precursor generation within the TCA cycle^75,76^. The citrate/α-KG ratio was approximately 2-to 6-fold higher in the retina than other ocular tissues, likely supporting the high biosynthetic demands of the retina for its rapid outer segment turnover^77^.

**Figure 6.**
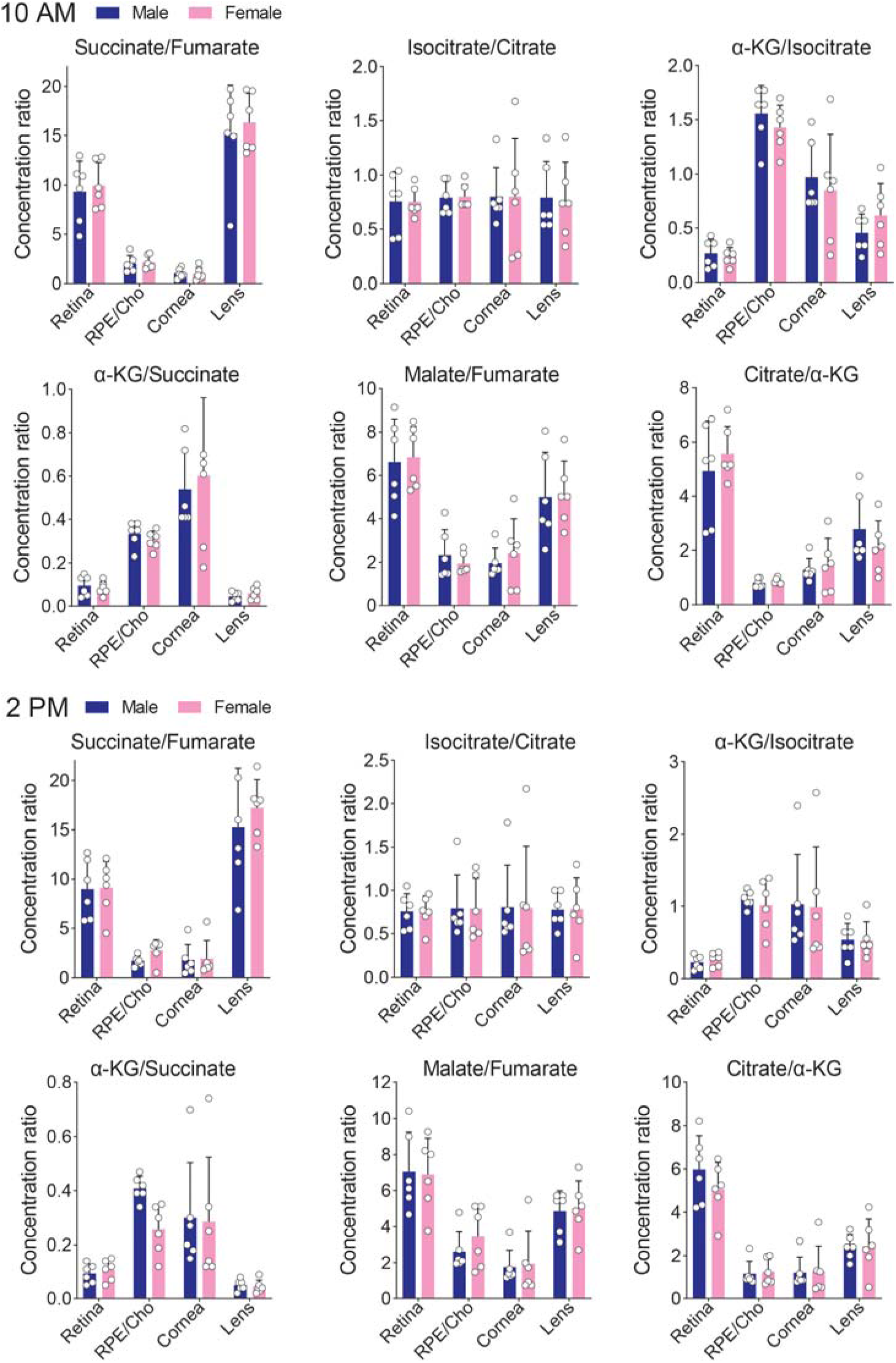
TCA metabolite ratios in male and female eye tissues at 10 AM and 2 PM. Ratios between key pairs of TCA cycle metabolites in the retina, RPE/choroid, cornea, and lens of male and female mice collected at 10 AM (A) and 2 PM (B). Metabolite ratios were calculated for each metabolite pair by dividing the mean value of the denominator metabolite (across all samples in the group) by the individual values of the numerator metabolite for each pair.

## 4. Discussion

In this study, we performed an absolute quantitation of TCA cycle metabolites, as well as pyruvate and lactate, across key ocular tissues to investigate metabolic differences between sexes and time points. We found strikingly different metabolic features in each ocular tissue (Figure 7): (1) the retina possesses the highest concentrations of most metabolites, consistent with strong bioenergetic and anabolic requirements; (2) RPE/choroid metabolites suggest large metabolic influxes and a highly reductive environment; and (3) cornea and lens metabolism are more sex-dependent, with the cornea possessing high malate to combat oxidative stress, while lens metabolites signify low metabolic activity and hypoxia. These findings reveal tissue-specific metabolism in ocular tissues, illustrating how TCA metabolism is geared toward supporting their unique functions.

**Figure 7.**
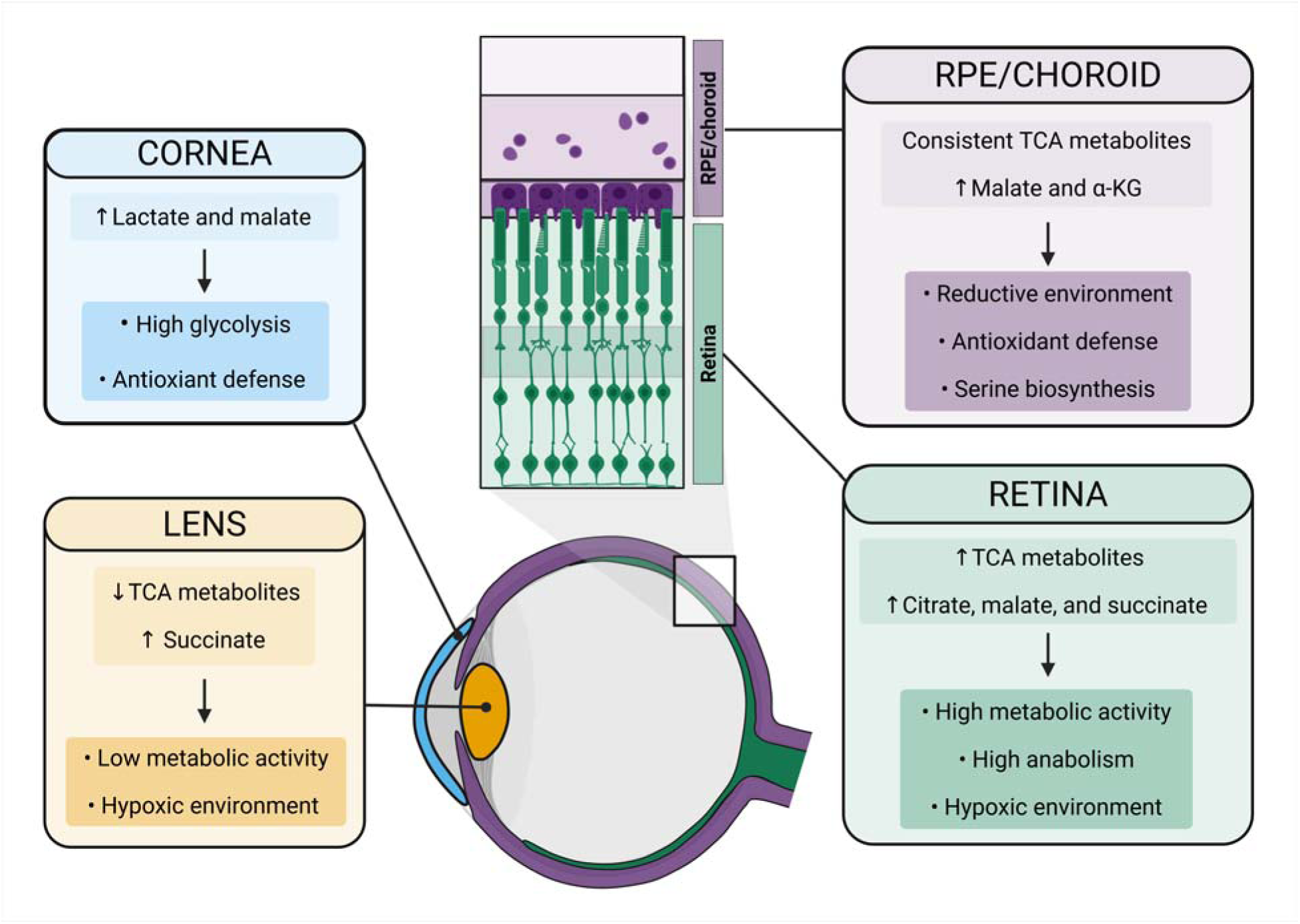
Summary of TCA cycle metabolite characteristics in different eye tissues. The retina (green), RPE/choroid (purple), cornea (blue), and lens (orange) show differences in TCA intermediate abundances. Key metabolites are up or downregulated in each tissue (shown in lighter boxes), which reflect or support their distinct environment or physiological requirements (shown in darker boxes).

### 4.1. Retina

Mitochondrial dysfunction in the retina is associated with many diseases, including Leber hereditary optic neuropathy, autosomal-dominant optic atrophy^78,79^, diabetic retinopathy^23,24,26^, retinitis pigmentosa^27–29^, AMD^25^, and glaucoma^23,30^. Our analysis further illustrates the retina’s dependence on mitochondrial metabolism, with concentrations of most TCA metabolites exceeding those in other ocular tissues by at least 10-fold. The retina contains high levels of malate, citrate, and succinate, strongly suggesting that these substrates are tuned to support the robust bioenergetic and biosynthetic demands of the retina^7,8^. Malate supports the MAS, enabling the shuttling of electrons between the mitochondria and cytoplasm^70^, which is important in energy-demanding tissues such as the heart, liver, and brain^80–82^. Indeed, whole-body and photoreceptor-specific disruption of the MAS causes impaired visual function and retinal degeneration, respectively^83,84^. Citrate accumulation is often a key indicator of an energy-rich and anabolic state^85,86^, and is typically abundant in tissues with high OXPHOS, including the retina^87–89^. Moreover, RPE cells use proline and glutamine to synthesize citrate that can be exported to the photoreceptors^15,72^, while photoreceptors employ citrate for fatty acid and phospholipid synthesis to support outer segment biosynthesis^90,91^.

One remarkable feature of the retinal mitochondrial intermediates was the high succinate/fumarate ratio. Succinate typically undergoes oxidation to fumarate, catalyzed by SDH^3,92^. Impaired respiration may cause the accumulation of reduced ubiquinone (ubiquinol), the electron acceptor for succinate oxidation, leading to a reverse SDH reaction in which fumarate is reduced to succinate^68,69^. Moreover, recent studies have shown that fumarate can act as a terminal electron acceptor in the mammalian ETC via the reversed SDH reaction, particularly in hypoxic tissues like the retina^68^. The high succinate/fumarate ratio also helps support metabolic cooperation, in which the oxygen-poor outer retina exports succinate to shuttle reducing power to the oxygen-rich RPE^16,93^.

### 4.2. RPE/choroid

Compared to other tissues, TCA metabolite levels in the RPE/choroid were remarkably balanced, although with relatively higher α-KG and malate. The RPE requires robust mitochondrial metabolism to process phagocytosed photoreceptor outer segments and nutrients from the choroidal circulation^15,94–96^. This active mitochondrial metabolism can create a highly reductive environment in the RPE/choroid, which may drive reductive carboxylation to generate citrate from α-KG, a highly active pathway in RPE cells^1,72,97,98^. This reductive carboxylation may help combat oxidative stress by promoting cytosolic NADPH generation through isocitrate dehydrogenase 1^72,99^. Malate can also support NADPH generation through malic enzyme ^100^. MAS activity may support serine biosynthesis by regenerating cytosolic NAD^+^, an essential cofactor for the rate-limiting enzyme of this pathway, phosphoglycerate dehydrogenase^101,102^. Serine is vital for RPE health and homeostasis, serving as a precursor for the synthesis of phospholipids, the antioxidant glutathione, and other amino acids^91,103,104^.

### 4.3. Cornea

The cornea had a high lactate concentration, which is consistent with early studies showing that the cornea converts approximately 85% of its glucose to lactate^33,34^. The crucial role of corneal mitochondrial metabolism is evident in inherited disorders such as Kearns-Sayre syndrome and Pearson syndrome, which result from large mitochondrial DNA deletions^105,106^. These conditions primarily affect the corneal endothelium ^53,107^, causing edema and a loss of transparency due to insufficient ATP production, which is needed to pump water out of the cornea^53,54,108–110^. Strikingly, malate is consistently the most abundant TCA metabolite in the cornea and is particularly high in female mice. In addition to its role in the MAS, malate is a key anaplerotic molecule that contributes to gluconeogenesis, transamination, and redox regulation for antioxidant defense^111,112^. The cornea is particularly vulnerable to oxidative stress, as free radicals are generated from incoming UV radiation and other sources^113^. Moreover, oxidative stress has been implicated in corneal diseases such as keratoconus^114^ and Fuchs endothelial corneal dystrophy^115^. Thus, the cornea may prioritize malate production as an antioxidant defense mechanism.

### 4.4. Lens

The lack of mitochondria in mature lens fibers makes the lens highly reliant on anaerobic glycolysis, but highly sensitive to oxidative stress, an important cause of cataracts^51,116–119^. To protect against oxidative stress, the lens maintains low oxygen tension, minimal mitochondrial activity, and high levels of antioxidants, and produces oxidation-resistant lipids and proteins^120–122^. Indeed, the lens had the lowest levels of TCA cycle metabolites, except for succinate, compared to all other tissues examined in this study. Although the epithelial cells overlying the anterior lens are rich in small mitochondria and primarily produce ATP through OXPHOS^47,48,50,123^, the anterior lens itself has a very low oxygen tension *in vivo* - about 10-fold lower than that of the cornea^118^. Given the limited oxygen supply, reverse SDH reactions in the lens are likely driven in a similar manner to the retina, leading to the reduction of fumarate to succinate^68,69^.

Both the cornea and lens demonstrated significant sex differences in TCA metabolites in the morning and afternoon. This may reflect the sex-specific differences in outer eye anatomy^60,124^, wound healing^125,126^, and pathologies such as cataracts^127,128^. These differences may arise due to sex hormone signaling; sex hormone receptors such as estrogen receptor alpha and androgen receptor are expressed in the lens and cornea^129–133^, while the circulating levels of these hormones vary widely between males and females^134^. Sex hormones, including 17β-estradiol, testosterone, and progesterone, are well-established regulators of both systemic and cellular metabolism^135–145^. The sex- and time point-dependent differences in the cornea and lens likely reflect the complex interplay between hormonal fluctuations and the circadian rhythm. Sex hormones fluctuate over a 24-hour cycle^146^, and these hormones are deeply connected to the body’s internal clock. For example, several core clock genes are regulated by estrogen response elements, and the brain and muscle ARNT-like 1-circadian locomotor output cycles kaput (BMAL1-CLOCK) complex drives rhythmic expression of estrogen receptor genes^147^. Our findings suggest that metabolic differences between sexes may underpin their varied susceptibility to ocular injury and disease.

## 5. Conclusions

Our findings demonstrate that each ocular tissue possesses a unique mitochondrial metabolic signature, and these tissue-specific differences may expose distinct vulnerabilities to disease. Our data also reveal a possible metabolic basis for sex differences in the outer eye, which should be further explored in future investigations.

## Abbreviations

α-KG: alpha-ketoglutarate
AMD: age-related macular degeneration
ECM: extracellular matrix
ETC: electron transport chain
GC-MS: gas chromatography-mass spectrometry
MAS: malate-aspartate shuttle
OXPHOS: oxidative phosphorylation
RPE: retinal pigment epithelium/epithelial
SDH: succinate dehydrogenase
TCA: tricarboxylic acid

## Acknowledgements

This work was supported by NIH Grants (EY026030, EY031324, EY032462), the Retina Research Foundation, NIH NIGMS P20GM144230 Visual Sciences COBRE grant to WVU, and an unrestricted challenge grant from Research to Prevent Blindness (RPB) to the Ophthalmology department at WVU.

## Conflicts of interest

The authors declare no conflicts of interest.

## Author contributions: CRediT

**Cloe Ratliff:** Formal analysis, investigation, data curation, writing - original draft, visualization, project administration. **David Hansman:** Data analysis, visualization, writing - original draft, writing - review & editing. **Yinxiao Xiang:** Investigation. **Artjola Puja:** Writing - original draft, visualization. **Mark Eminhizer:** Investigation. **Tuan Ngo:** Investigation. **Jinyu Lu:** Visualization. **Isabella Mascari:** Data analysis and visualization. **Diana Alabdallat:** Writing - review & editing. **Jianhai Du:** Supervision, project administration, funding acquisition, writing - review & editing, methodology, conceptualization.

## Notes

### Competing Interest Statement

The authors have declared no competing interest.

