## Supplementary material for "Absolute quantification of TCA cycle intermediates in mouse ocular tissues reveals distinct tissue- and sex-specific mitochondrial metabolism": Supplementary figures 1-4.docx

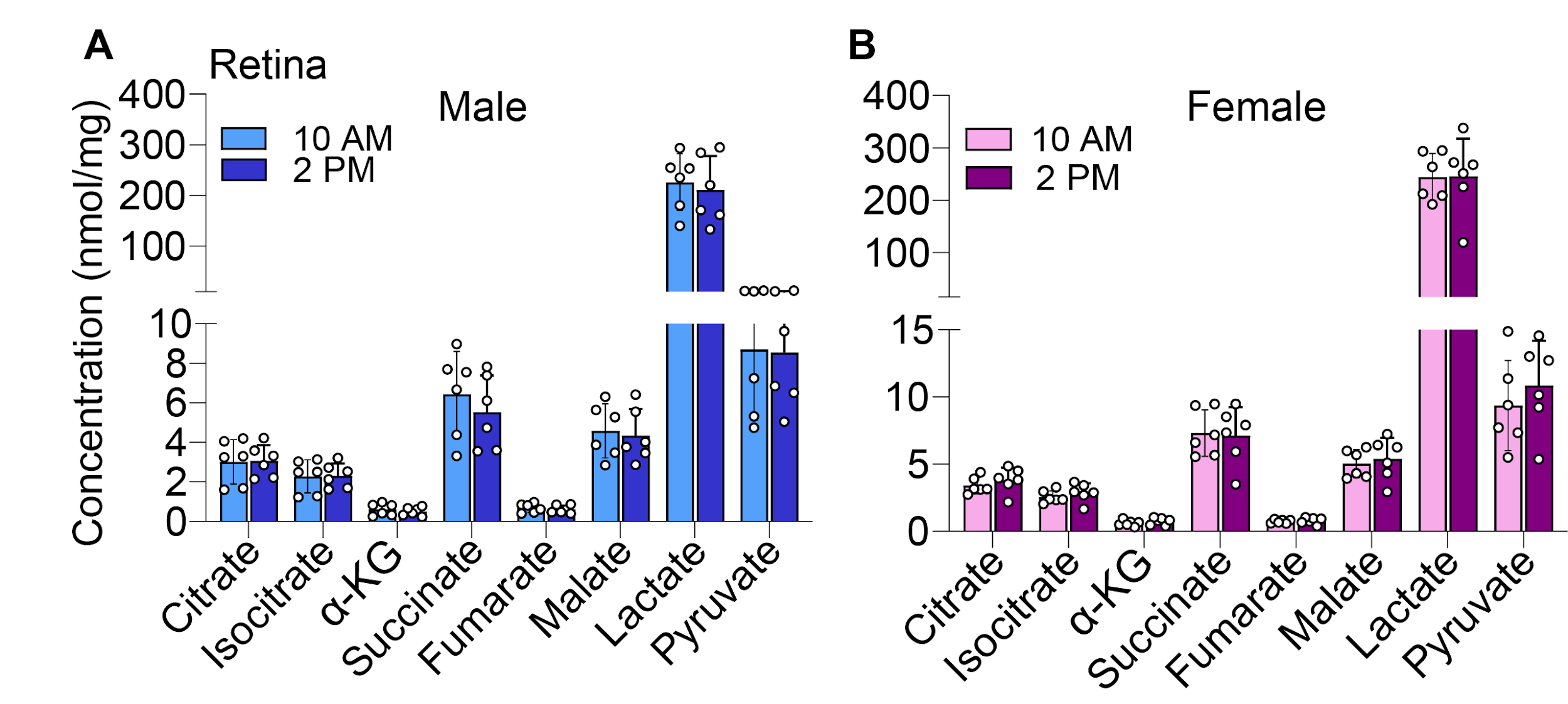

**Figure S1. Comparison of retinal TCA cycle metabolite abundance between 10 AM and 2 PM.** Lactate, pyruvate, and TCA metabolite concentrations in male (A) and female (B) mouse retinas collected at 10 AM and 2 PM. No statistical significance (p<0.05) between samples collected at 10 AM and 2 PM were detected using an unpaired, two-tailed Student’s t-test. α-KG = α-Ketoglutarate.

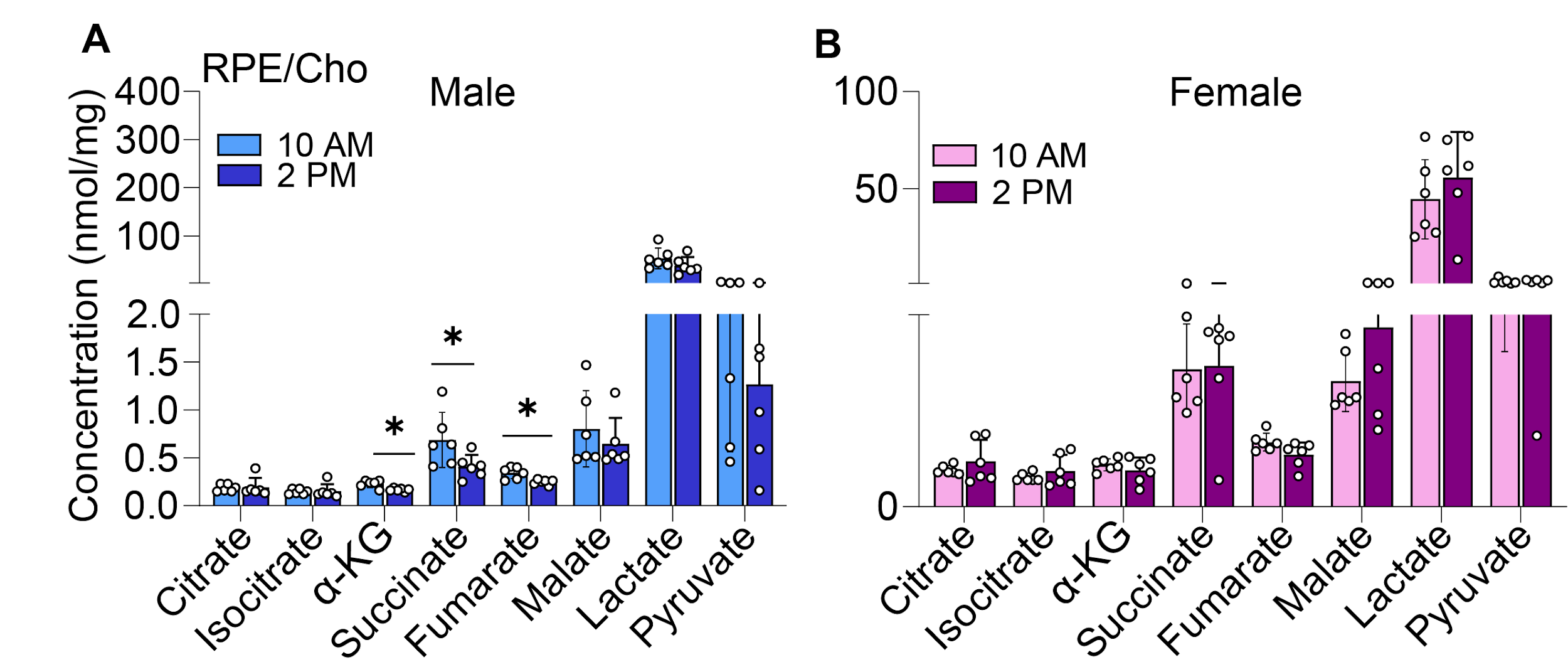

**Figure S2. Comparison of RPE/choroid TCA cycle metabolite abundance between 10 AM and 2 PM.** Lactate, pyruvate, and TCA metabolite concentrations in male (A) and female (B) mouse RPE/choroids collected at 10 AM and 2 PM. *P<0.05 between samples collected at 10 AM vs. 2 PM, detected using an unpaired, two-tailed Student’s t-test. α-KG = α-Ketoglutarate.

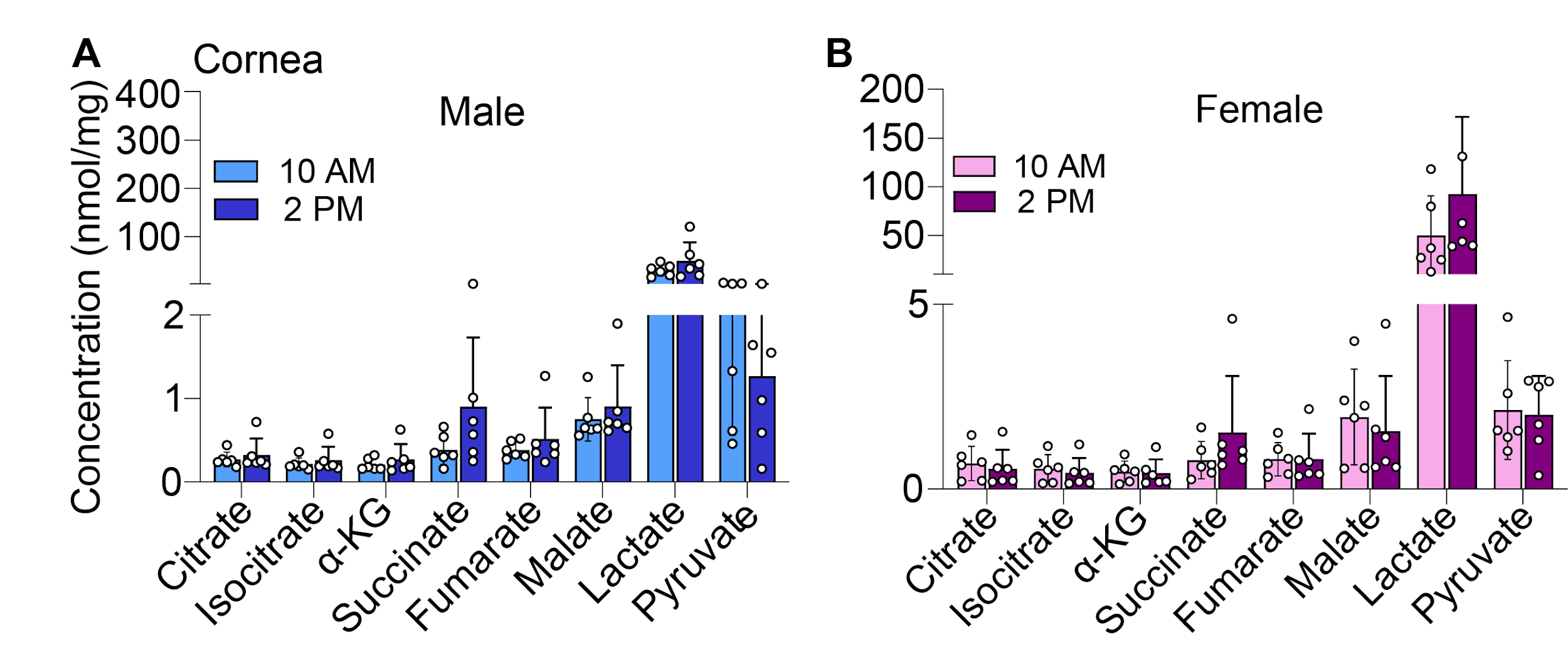

**Figure S3. Comparison of corneal TCA cycle metabolite abundance between 10 AM and 2 PM.** Lactate, pyruvate, and TCA metabolite concentrations in male (A) and female (B) mouse corneas collected at 10 AM and 2 PM. *P<0.05 between samples collected at 10 AM vs. 2 PM, detected using an unpaired, two-tailed Student’s t-test. α-KG = α-Ketoglutarate.

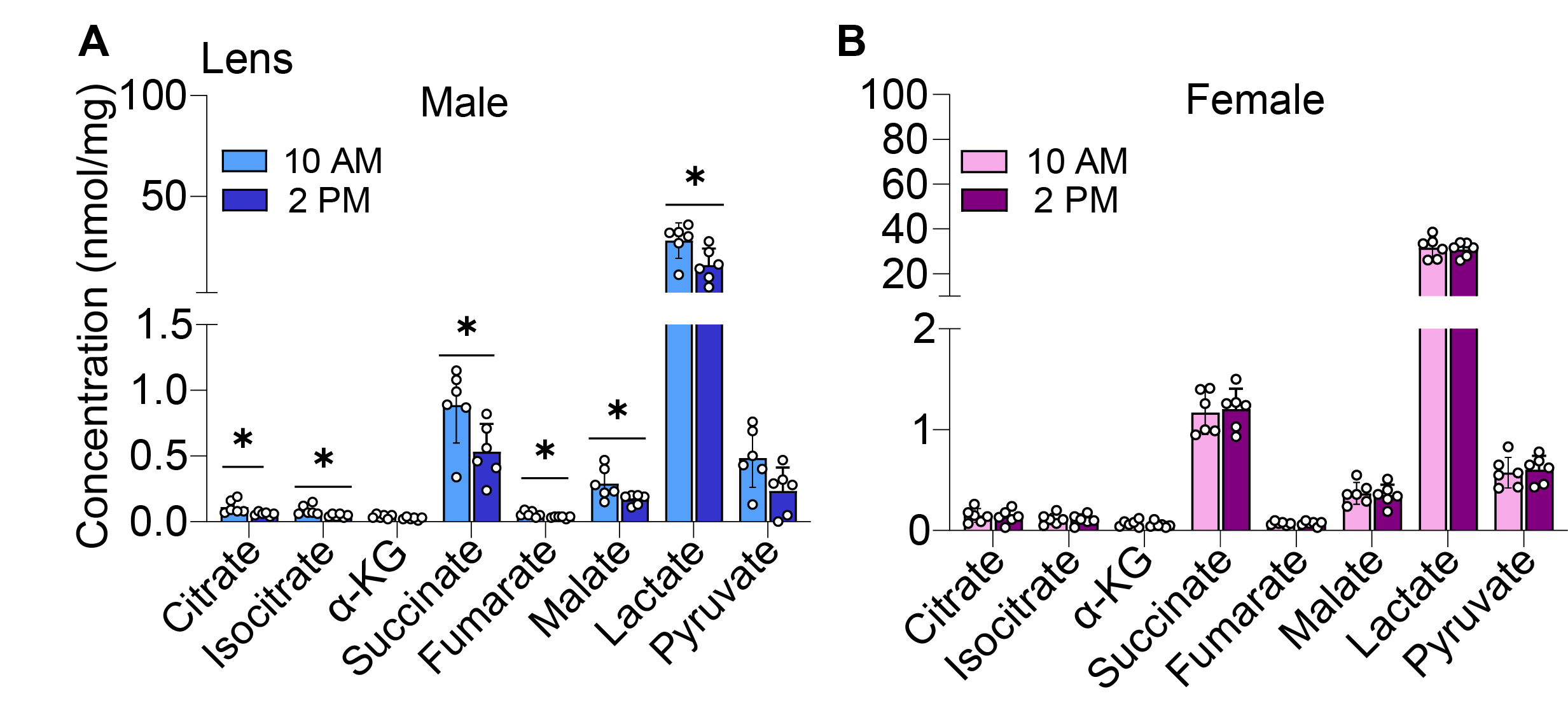

**Figure S4. Comparison of lens TCA cycle metabolite abundance between 10 AM and 2 PM.** Lactate, pyruvate, and TCA metabolite concentrations in male (A) and female (B) mouse lenses collected at 10 AM and 2 PM. *P<0.05 between samples collected at 10 AM vs. 2 PM, detected using an unpaired, two-tailed Student’s t-test. α-KG = α-Ketoglutarate.
