## Supplementary material for "Absolute quantification of TCA cycle intermediates in mouse ocular tissues reveals distinct tissue- and sex-specific mitochondrial metabolism": Table S1 Key Resources.docx

**Absolute quantification of TCA cycle intermediates in mouse ocular tissues reveals distinct tissue- and sex-specific mitochondrial metabolism**

Cloe Ratliff^1,2*^, David Hansman^1,2*^, Tuan Ngo^1,2^, Yinxiao Xiang^1,2^, Artjola Puja^1,2^, Mark Eminhizer^1,2^, Jinyu Lu^1,2^, Isabella Mascari^1,2^, Diana Alabdallat^1,2^, Jianhai Du^1,2^*

| **Mass Spectrometry (GC-MS)** | **Source** | **Identifier** |
| --- | --- | --- |
| Water, Optima™ LC/MS Grade | Fisher Chemical | 166415 |
| Methanol | Fisher Chemical | 164905 |
| Methoxyamine hydrochloride | Sigma-Aldrich | 226904 |
| Pyridine | Sigma-Aldrich | 270970 |
| N-tert-Butyldimethylsilyl-N-methyltrifluoroacetamide | Sigma-Aldrich | 394882 |
| DB-5ms GC Column, 30 m, 0.25 mm, 0.25 um | Agilent Technologies | 1225532 |
| Gel Pump | Savant | GP110 |
| Speed vac Plus | Savant | SC110A |
| Centrifuge | Eppendorf | 5424 |
| **Animals** |  |  |
| C57BL/6J | Jackson Lab | 000664 |
| **Protein Normalization** |  |  |
| Pierce™ Dilution-Free™ Rapid Gold BCA Protein Assay | Thermo Fisher | A55860 |
| Gibco™ Phosphated-Buffer Saline (PBS), pH 7.4 | Fisher Chemical | 10-010-031 |
| Bovine Serum Albumin (BSA) | Sigma-Aldrich | A7030 |
